# High-throughput, real-time monitoring of engineered skeletal muscle function using magnetic sensing

**DOI:** 10.1101/2022.05.20.492879

**Authors:** Alec S.T. Smith, Shawn M. Luttrell, Jean-Baptiste Dupont, Kevin Gray, Daniel Lih, Jacob W. Fleming, Nathan J. Cunningham, Sofia Jepson, Jennifer Hesson, Julie Mathieu, Lisa Maves, Bonnie J. Berry, Elliot C. Fisher, Nathan J. Sniadecki, Nicholas A. Geisse, David L. Mack

## Abstract

Engineered muscle tissues represent powerful tools for examining tissue level contractile properties of skeletal muscle. However, limitations in the throughput associated with standard analysis methods limit their utility for longitudinal study, high throughput drug screens, and disease modeling. Here we present a method for integrating 3D engineered skeletal muscles with a magnetic sensing system to facilitate non-invasive, longitudinal analysis of developing contraction kinetics. Using this platform, we show that engineered skeletal muscle tissues derived from both induced pluripotent stem cell and primary sources undergo improvements in contractile output over time in culture. We demonstrate how magnetic sensing of contractility can be employed for simultaneous assessment of multiple tissues subjected to different doses of known skeletal muscle inotropes as well as the stratification of healthy versus diseased functional profiles in normal and dystrophic muscle cells. Based on these data, this combined culture system and magnet-based contractility platform greatly broadens the potential for 3D engineered skeletal muscle tissues to impact the translation of novel therapies from the lab to the clinic.

## Introduction

Three-dimensional engineered muscle tissues (EMTs) are a well-established method for studying the contractile output of skeletal muscle *in vitro* ^1-22^. The maintenance of muscle cells within a 3D environment better recapitulates the cell-cell and cell-matrix interactions that are critical for normal muscle development and helps promote a more mature *in vitro* phenotype for downstream analysis ^3, 9, 10^. Despite their relative ubiquity in small-scale academic studies, integration of EMT technologies into large-scale drug screens and commercial workflows necessitates a scalable approach to functional analysis. Typically, EMT function is determined either by optical measurement of contraction-induced substrate deformation ^20, 21^ or direct quantification of force output via connection of the tissue to a force transducer ^10, 16, 22^. Such methods often limit analysis to a single tissue at a time and demand a high degree of operator skill and/or time to achieve actionable data. While some aspects of microscopy-based, substrate-deflection measurements can be automated, they require dedicated microscopy equipment and custom image analysis routines to ensure accurate and consistent results. Furthermore, such technologies still prevent simultaneous assessment of multiple tissues, making collection of dose response data challenging.

Previous work has demonstrated that magnetic field sensing can be employed as a means to assess engineered cardiac tissue function in parallel across multiple constructs ^23^. Engineered tissues were suspended between one rigid and one flexible post in a 24-well lattice. A magnet was embedded in the tip of each flexible post and the lattice positioned above giant magnetoresistive (GMR) sensors. In this system, tissue contraction caused the flexible posts to deflect, moving the position of the embedded magnets, which in turn produced a voltage change at the GMR sensor that could be used to measure force and frequency. Here, we describe the adaptation of this technology to facilitate analysis of engineered skeletal muscle tissues derived from both human induced pluripotent stem cells (iPSCs) and primary sources. We demonstrate the ability for these constructs to undergo functional maturation over time, respond to chemical challenge, and to stratify healthy and diseased phenotypes. We provide workflows for the use of this technology in performing dose-response drug studies and characterize the morphological development of the cells maintained in this system. In so doing, we describe a holistic approach to skeletal muscle modeling *in vitro*, including methods to produce highly contractile muscle from multiple sources, integration and further differentiation of these cells within 3D engineered constructs, and subsequent, longitudinal analysis of functional output in a non-invasive and highly scalable manner that enables higher-throughput screening modalities.

## Materials & Methods

### Maintenance of hiPSC lines

We utilized urine-derived iPSC lines generated internally by our group throughout this study. The UC3-4 and UCS-2 lines were derived using similar methods ^24^. However, while the UC3-4 line was generated using first-generation lentiviruses, the UCS-2 line was created using footprint-free, non-integrating Sendai viruses (2.0 Sendai Reprogramming Kit; Thermo Fisher Scientific, Waltham, MA, USA), therefore representing an advancement on the previously published line. Both UC3-4 and UCS-2 hiPSCs were frozen in mFreSR medium (Stem Cell Technologies, Vancouver, Canada) and stored under cryogenic conditions in a vapor phase liquid nitrogen dewar. On the day of plating, vials of cells were removed from liquid nitrogen storage, thawed, pelleted in a tabletop centrifuge run at 300 g for 3 minutes, and resuspended in mTeSR medium (Stem Cell Technologies) supplemented with 10 µM Y-27632 (Thermo Fisher Scientific), a specific inhibitor of Rho kinase activity. Cells were then plated on surfaces that had previously been coated overnight with Matrigel (Thermo Fisher Scientific) diluted 1 to 60 in DMEM/F12 medium (Thermo Fisher Scientific). Y-27632 was removed from the culture medium the first day after plating and cells were then fed daily with fresh mTeSR. Cells were incubated at 37°C/ 5% CO_2_ until iPSC colonies filled the field of view when visualized using an Eclipse TS100 microscope (Nikon, Tokyo, Japan) fitted with a 10X lens. At this point, cells were lifted off of the culture surface using TrypLE Select (Thermo Fisher Scientific), collected, spun down, and resuspended in fresh mTeSR with mild trituration to gently break up cell clusters before being split across the desired number of Matrigel-coated plates. During continued culture, any iPSC colonies displaying irregular boundaries, significant space between cells or low nuclear to cytoplasmic ratios were carefully marked and removed from culture using a fire-polished, sterile glass pipette.

### Generation of dystrophin-null hiPSCs

Working with the University of Washington’s Institute for Stem Cell and Regenerative Medicine Ellison Stem Cell Core, dystrophin-null genotypes were engineered into the UC3-4 wild type line using methods described previously ^25, 26^. Cas9 guide RNAs (gRNA) targeting the DMD locus were designed using CRISPoR. Three gRNA were chosen, targeting intron 44 (AAAAACTGGAGCTAACCGAG), intron 45 (ATATACTTGTGGCTAGTTAG) and exon 54 (TGGCCAAAGACCTCCGCCAG) (**Figure S1A**). To limit the exposure of the DNA to the genome editing enzyme and reduce off-target events, the Cas9 protein and guide RNAs were introduced as ribonucleoprotein (RNP) complexes. One million UC3-4 iPSCs were electroporated with Cas9 (0.6uM, Sigma) and gRNA (1.5uM, Synthego) using Amaxa nucleofector (Human Stem Cell kit 2) in presence of ROCK inhibitor. Individual clones were handpicked and plated into 96 well plates. DNA was extracted using Quick Extract DNA extraction solution (Epicentre#QE09050) and nested PCR was performed using Forward (CATGGAACATCCTTGTGGGGA) and Reverse (CTGAGATTCAGCTTGTTGAAGTTA) primers for exon 45 region and Forward (AGTTTGTCCTGAAAGGTGGGT) and Reverse (CCTCCCAAGCTCCAGTTTAGC) primers for exon 54 region. The PCR products were purified using EXO-SAP enzyme (ThermoFisher) and sent for Sanger sequencing analysis (through Genewiz). Clones harboring the correct DMD mutation were amplified and sent for G-band karyotyping (outsourced to Diagnostic Cytogenetics Inc., Seattle, WA, USA. Data not shown). The collected sequencing data confirmed the presence of a dystrophin-null phenotype in our engineered DMD line (**Figure S1B**). Once established, mutant lines derived from single clones were expanded and maintained as described above.

### Differentiation, purification, and expansion of hiPSC-derived skeletal muscle myoblasts

Differentiation of hiPSCs and commitment to the myogenic lineage was achieved using a protocol modified from those published previously ^27, 28^ (**Figure 1A**). First, UCS-2 hiPSCs were plated at 15,000 cells/ cm^2^ and grown to roughly 40% confluency in mTeSR. At this point, the medium was replaced with a differentiation medium consisting of DMEM/F12 basal medium, 1x non-essential amino acid supplement (Thermo Fisher Scientific), 1x Insulin-Transferrin-Selenium supplement (Thermo Fisher Scientific), 3 µM CHIR99021 (Axon Medchem, Reston, VA, USA), and 0.2 µg/mL LDN193189 (Miltenyi Biotec, Waltham, MA, USA). On day 3 post induction, this medium was further supplemented with 20 ng/mL bFGF (R&D Systems, Minneapolis, MN, USA). On day 6 post induction, the CHIR99021 and the Insulin-Tranferrin-Selenium supplement were removed and replaced with 15% knockout serum replacement (KSR; Thermo Fisher Scientific), 2 ng/ mL IGF-1 (R&D Systems), and 10 ng/ mL HGF (R&D Systems). On day 8 post induction, bFGF, HGF, and LDN193189 were all removed from the differentiation medium. On day 12 post induction, HGF was again supplemented into medium and this composition was then maintained until day 28. Throughout this period, medium was changed daily.

**Figure 1:**
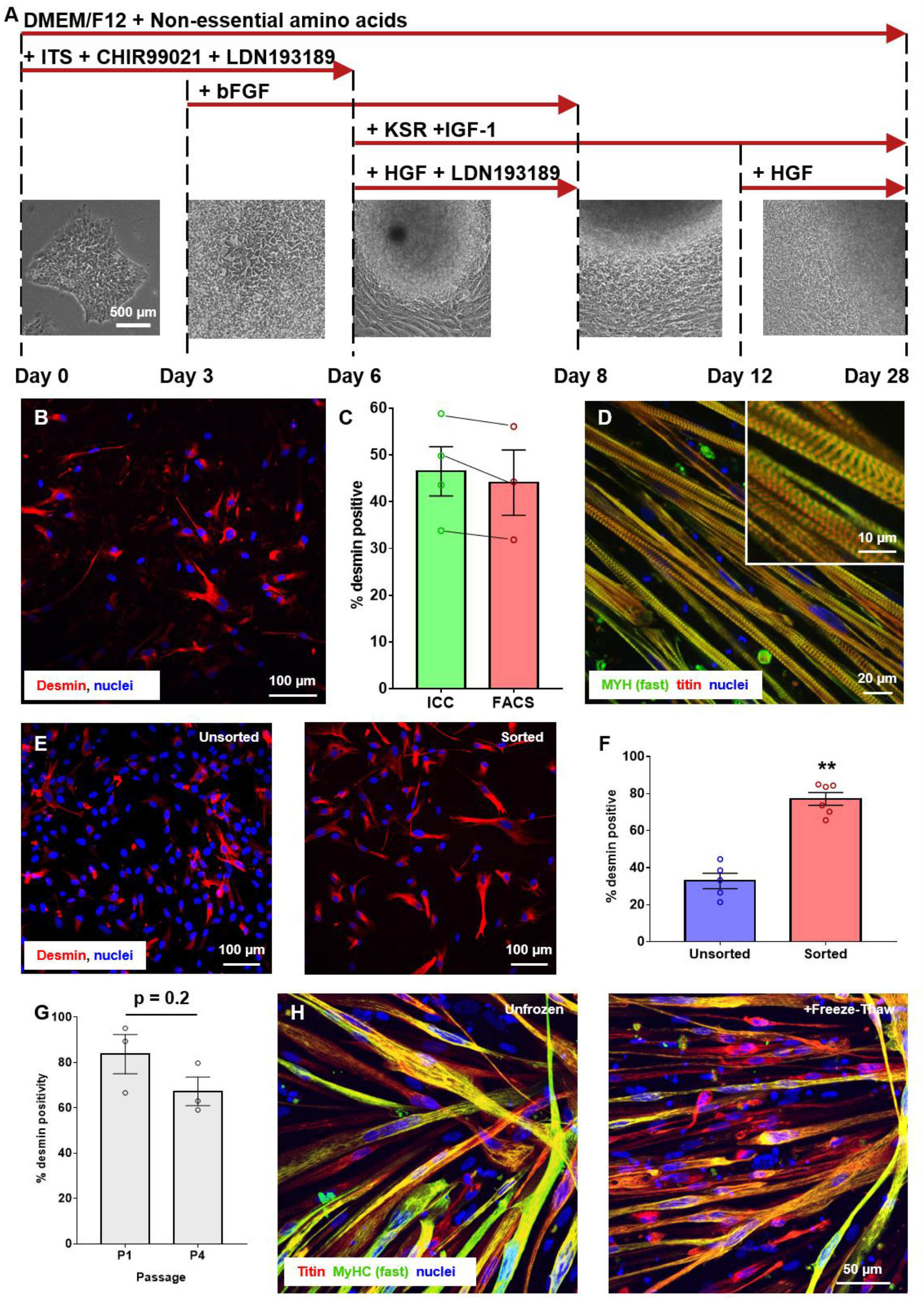
Human iPSC-derived skeletal muscle production and characterization. (A) Schematic illustration of the differentiation protocol employed in this study. Inset images detail representative images of cells at different timepoints throughout this culture. (B) Day 28 iPSC-derived myoblasts stained for expression of the muscle specific intermediary filament, desmin. (C) Quantification of desmin purity in iPSC-derived myogenic cultures at day 28 post-induction by immunocytochemistry (ICC) and fluorescence activated cell sorting (FACS). Black lines link data points collected from the same differentiation run. n = 4. (D) Representative image of day 35 myogenic cultures subjected to serum-starvation to promote the formation of multinuclear myotubes. Cells were stained with an antibody against m-line titin to illustrate the development of sarcomeric structures within these cultures. (E) Representative images of myogenic cultures pre- and post-live capture FACS. (F) Quantification of desmin purity in pre- and post-FACS myogenic populations. **p < 0.01, n = 6 per group. (G) Quantification of desmin purity at passage 1 (P1) versus passage 4 (P4). No statistically significant difference in mean purity was detected between groups, n = 3. (H) Representative immunostained images of iPSC-derived myotubes produced from populations never subjected to cryogenic storage and those that underwent a single freeze-thaw event.

At day 28 post induction, confluent cultures were passaged using TrypLE Select and replated on Matrigel-coated surfaces for further expansion. Cells at this stage stained positive for the muscle-specific marker desmin (**Figure 1B,C**) and could be induced to fuse, forming multinuclear myotubes with distinct sarcomeric patterning, via serum starvation (**Figure 1D**). All subsequent expansion steps were performed using a commercially-sourced skeletal muscle growth medium (Lonza Group AG, Basel Switzerland) and cells were fed every 2 to 3 days. At day 32 post-induction, differentiated cells were again lifted off their culture surfaces using TrypLE Select and subjected to fluorescence activated cell sorting (FACS) after being stained with an erbB3 antibody (Biolegend, San Diego, CA, USA) conjugated to phycoerythrin. FACS purification was performed with the support of the Cell Analysis Facility in the Department of Immunology at the University of Washington. Previous studies demonstrated that erbB3 FACS can be used to purify myogenic precursors from non-myogenic contaminants ^29^ and was employed here as a means to increase the purity of our myogenic cultures (**Figure 1E,F**). Purified cultures were further expanded on Matrigel-coated plates to generate sufficiently large populations of cells for downstream experiments. All cells used in this study were isolated from passage 1 to 4 cells as analysis of desmin positivity indicated the persistence of a relatively stable myogenic fraction in these cultures through this level of expansion (**Figure 1G**). Expanded iPSC-derived myoblasts could then be cryogenically-stored using mFreSR medium (Stem Cell Technologies) until needed. Frozen and then thawed cells retained their capacity to fuse under serum starvation conditions, producing multinuclear myotubes similar to those obtained from populations that had never been subjected to cryogenic storage (**Figure 1H**).

### Culture of primary human myoblasts

Primary human myoblasts obtained from biopsy of healthy adult donors were sourced commercially (Lonza, CC-2580) and maintained in culture according to the vendor’s protocols. Vials of cells from the vendor were thawed and plated on Matrigel coated tissue culture surfaces and expanded in Lonza’s skeletal muscle growth medium (SkGM-2). Expanded cells were frozen in mFreSR medium (Stem Cell Technologies) until required for downstream experimentation. All experiments detailed in this manuscript were derived from cells isolated between passages 1 to 4.

### Generation of 3D engineered muscle tissues (EMTs)

EMTs were established using Curi Bio’s Magnetometric Analyzer for eNgineered Tissue ARRAY (Mantarray) casting platform (**Figure 2A**), and according to the manufacturer’s protocols. Briefly, muscle cells were enzymatically-dissociated and lifted from their culture surfaces, counted, and spun down at 300 g for 5 minutes. 750,000 cells were then resuspended in 107 µL Lonza growth medium, supplemented with 30 µL Matrigel, 10 µL 50 mg/ mL fibrinogen (Sigma-Aldrich), and 3 µL 100 U/ mL thrombin (Sigma-Aldrich) to create a total seeding volume of 150 µL and a seeding density of 5 million cells/ mL. This cell suspension was applied to Mantarray casting wells housing the two-post array and incubated at 37°C/ 5% CO_2_ for 80 minutes to facilitate hydrogel formation. Following incubation, 1 mL of Lonza growth medium was added to the casting wells to aid separation of the gel from the substrate. The two-post array supporting the newly formed hydrogels was then lifted out of the casting plate and transferred to a fresh 24-well plate containing Lonza growth medium supplemented with 2 g/L aminocaproic acid (Sigma-Aldrich) to inhibit fibrinolysis. One day after plating, EMTs were transferred to a low-serum (2%) medium supplemented with 2 g/L aminocaproic acid (ACA) to induce myoblast fusion. Cultures were maintained in the low-serum medium for the remainder of the culture period and were fed with fresh medium every 2 to 3 days from this point onward. ACA concentration was raised to 5 g/L between days 7 and 10 to combat matrix degradation in long-term cultures. Primary constructs were maintained in 5 g/L ACA throughout the entire culture period.

**Figure 2:**
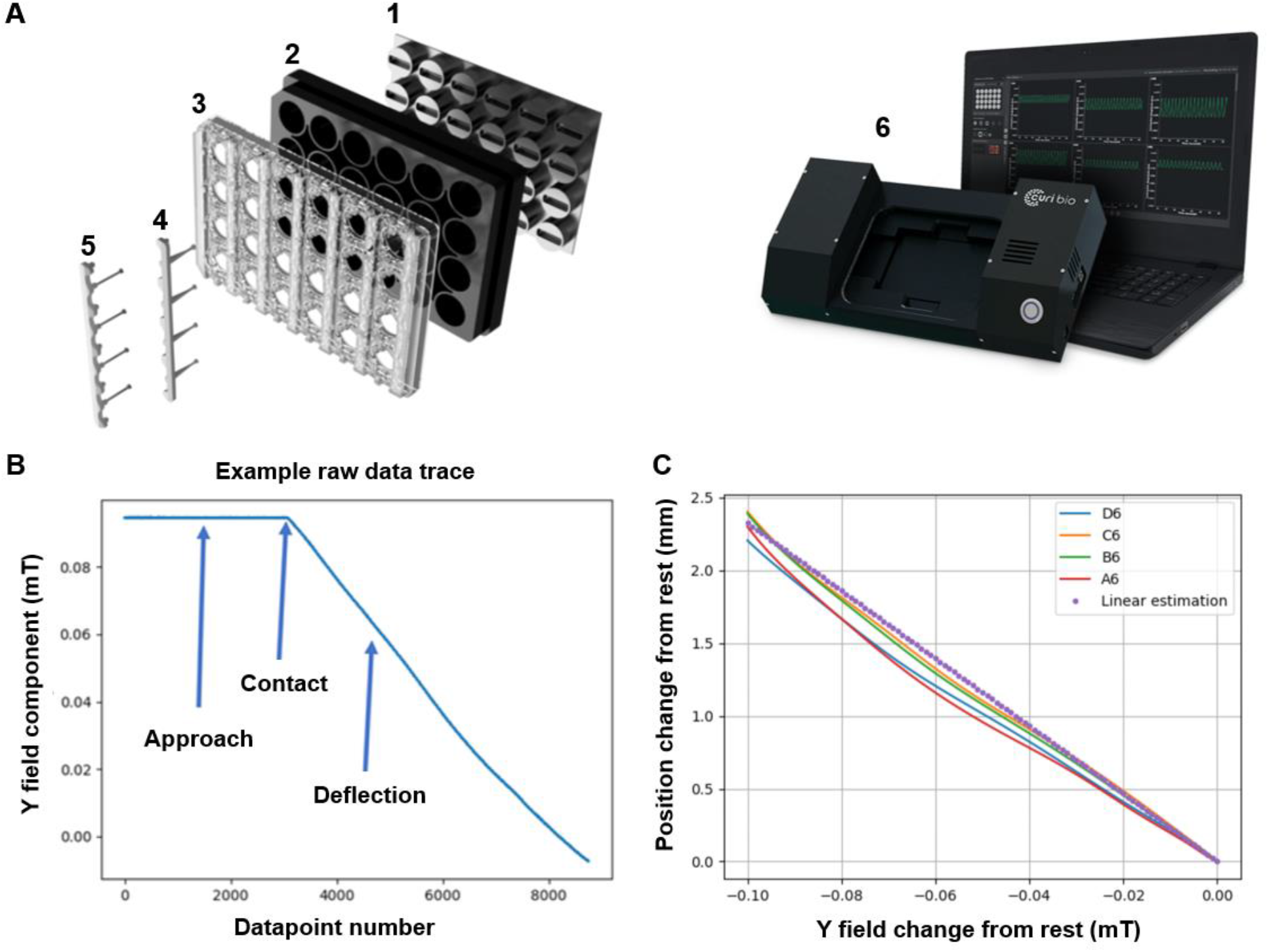
Calibration of the Mantarray device. (A) Exploded diagram of the Mantarray plate. Tissues are cast in the casting plate (1) and then transferred to the culture plate (2). The plate lattice (3) supports both the rigid post (4) and the flexible post containing the embedded magnet (5). Post deflections are measured by mounting the culture plate onto the Mantarray hardware (6). (B) Example raw data trace illustrating the change in magnetic field deflection as the probe approaches and then deflects the flexible post within a single well of the Mantarray plate. (C) Recorded movements from analysis of 4 wells normalized for variability in post starting position.

### Mantarray calibration

The Mantarray hardware contains an electronic printed circuit board that supports an array of fluxgate sensors and their ancillary electronic components required for the processing and digitization of the sensors’ output. The EMT plate is placed within this hardware so that the post tips are a few millimeters from the fluxgate sensor. As the tissue contracts, the deformable post and its embedded magnet move in response within the magnetic field of the sensor. As the magnetic field at the sensor changes in response to contraction, the fluxgate device outputs a voltage change. The change in detected voltage is linear within the normal range of movement for the flexible posts and does not change over the life of the device. Calibration was achieved via a custom-designed micro-positioner connected to the flexible post on the EMT plate. The micro-positioner actuated the post at a controlled rate across a known distance, thereby moving the magnet it from its resting position (**Figure 2B**). Using this method, it was possible to correlate the recorded field at a given time point with respect to the distance that the post had been deflected (**Figure 2C**). This process was repeated for 4 posts, with the lattice holding the posts removed and replaced prior to each repetition, as is likely to occur during regular use. Once calibrated, magnetically-induced voltage changes recorded in response to EMT contraction could be converted to physical movement, in μm or mm. Then, using Euler-Bernoulli beam theory, modeling the post as a uniform cantilever with a point load at its end, the mechanical properties of the fabricated post could be used to quantify contraction force. Calibration calculations are performed internally by the Mantarray software suite and so require no expertise on the part of the operator.

### Measurement of EMT contraction

EMT contraction was elicited via broad-field electrical stimulation. A purpose-built plate lid supporting graphite electrodes that was compatible with the Mantarray hardware was used in conjunction with a C-Pace EP in-culture stimulator (IonOptix, Westwood, MA, USA). EMT cultures were subjected to electrical pulses of 10 V for 5 ms at a rate of 1 Hz for analysis of twitch contractions and 10 to 30 Hz to analyze tetanic responses. Post-deflection was measured by tracking the deformation of the magnetic field caused by movement of the magnet embedded in the flexible post using the dedicated Mantarray software suite. As the described system is non-invasive, individual EMTs could be monitored for changes in function over time. For the purposes of these studies, contractile properties of cultured EMTs were measured every 3 to 4 days between days 7 and 14 post tissue formation. However, the system allows for more regular measurements (daily, hourly, etc.) if required by the end user.

### Drug treatment and analysis of EMT functional response

Ryanodine was dissolved in dimethylsulphoxide (DMSO; Med Chem Express, Monmouth Junction, NJ, USA) to create a 1000X stock solution. The drug was then diluted in culture medium so that the final DMSO concentration exposed to cells was 0.1%. 2,3-Butanedione monoxime (BDM) was dissolved in media to make a 50 mM stock solution. Drug solutions were incubated with EMTs at 37°C/ 5% CO_2_ for 5 minutes prior to recording. For each drug analyzed, DMSO and media-only control samples were recorded and all drug-induced changes were compared to levels observed in these controls. For washout studies, EMTs were washed twice with PBS and returned to maintenance medium for 1 hour before rerecording contractile performance on the Mantarray hardware.

### Immunofluorescent staining

EMTs were fixed in 4% paraformaldehyde for 18-24 hours at 4°C. Whole EMTs were then dehydrated through an ethanol series (70%, 80%, 95%, 95%, 100%, 100%, for 30 minutes each) at room temperature. Dehydrated EMTs were moved through 3 changes of 100% xylene, for 30 minutes each at room temperature, and then transferred to a 1:1 mixture of xylene and paraffin wax for 30 minutes at 60°C. EMTs were subsequently moved through 3 changes of 100% paraffin wax, each for 30 minutes at 60°C, before being embedded in fresh paraffin wax. Next, 7 µm sections were cut on a Leica RM 2135 rotary microtome and mounted on Superfrost Plus microscope slides (Fisher Scientific).

Slides were incubated at 60°C for 10 minutes and moved through 3 changes of 100% xylene, each for 10 minutes at room temperature. Slides were then rehydrated through an ethanol series (100%, 100% for 5 minutes each, followed by 95%, 95%, 80%, 70%, for 3 minutes each) and rinsed in ddH20. Antigen retrieval was performed by incubating slides in 1X citrate buffer (Sigma-Aldrich) at 95°C for 10 min. Slides were allowed to cool in the citrate buffer on the benchtop for 30 minutes or until the final temperature of the buffer was approximately 50-60°C. Sections were blocked in 0.2% Triton X-100 + 1% BSA in PBS (blocking buffer) for 30 minutes at room temperature. Primary antibodies were diluted in blocking buffer and sections were incubated overnight at 4°C. Control sections were incubated overnight in blocking buffer without primary antibodies. The following day, sections were washed 3 times for 10 minutes each in PBS. Secondary antibodies were diluted in PBS and sections were incubated for 1 hour at room temperature with 4’,6-diamidino-2-phenylindole (DAPI; Sigma-Aldrich) for nuclei visualization. Slides were then washed 3 times for 10 minutes each in PBS and coverslips were applied with Vectashield antifade mounting medium (Vector Laboratories).

Cells cultured in 2D on glass coverslips were fixed in paraformaldehyde (4%; Thermo Fisher Scientific) for 15 minutes. Samples were then permeabilized and blocked with Triton-X-100 (0.2%; Sigma-Aldrich) and goat serum (5%; Thermo Fisher Scientific) in PBS for 1 hour at room temperature. Following blocking, samples were incubated with primary antibodies diluted in goat serum (1%) in PBS overnight at 4°C. The next day, cells were washed 3 times with PBS. They were then incubated at room temperature for 2 hours in a secondary antibody solution containing fluorophore-conjugated, secondary antibodies diluted in goat serum (1%) in PBS. Coverslips were mounted onto glass microscope slides using Vectashield containing DAPI for visualization of cell nuclei.

Images were taken at the University of Washington’s Garvey Imaging Core using a Nikon A1 Confocal System on a Ti-E inverted microscope platform. 12-bit 1024×1024 pixel images were acquired with Nikon NIS Elements 3.1 software. Antibodies used in this study were as follows: rabbit anti-desmin (1 in 200, Abcam), mouse anti-myosin heavy chain (1 in 500, Sigma-Aldrich), rabbit anti-m-line titin (1 in 300, Myomedix), mouse anti-MF20 (1 in 200, Developmental Studies Hybridoma Bank, Iowa City, Iowa, USA), rabbit polyclonal anti-dystrophin (1 in 1000, Abcam), Alexafluor-594 conjugated goat-anti-rabbit secondary antibody (1 in 200, ThermoFisher Scientific), Alexafluor-568 conjugated donkey-anti-rabbit secondary antibody (1 in 500, ThermoFisher Scientific), Alexafluor-488 conjugated donkey-anti-mouse secondary antibody (1 in 500, ThermoFisher Scientific), and Alexafluor-488 conjugated goat-anti-mouse secondary antibody (1 in 200, ThermoFisher Scientific). Analysis of stained as well as bright-field images of cultured cells were performed using ImageJ with standard plugins.

### Statistical analysis

All experiments were performed with technical duplicates or triplicates, and repeated using at least 3 independently-differentiated iPSC skeletal muscle populations. Significant differences between groups were evaluated using either unpaired, two-tailed t-tests or Mann Whiney-U tests depending on whether the data was normally distributed. One-way analysis of variance (ANOVA), or ANOVA on ranks depending on data normality, with post hoc tests for multiple comparisons was used to compare data sets with more than two experimental groups. All drug studies were performed across at least 5 doses and the data was then used to plot dose response curves for calculation of EC_50_ values from each metric analyzed. Dose response curves were fit to the collected data by running a nonlinear regression (log(agonist/antagonist) versus normalized response) with a least-squares fit in GraphPad Prism. In each case, dose response curves were normalized by expressing untreated values as 0% change, and the % change observed in treated samples at the highest dose tested as 100% change. In all experiments, a p value of less than 0.05 was considered significant. All statistical tests were performed using the GraphPad Prism statistics software package (GraphPad Software Inc.).

## Results

### Characterization of iPSC-based EMT structure and function on Mantarray

Human UCS-2 iPSC-derived EMTs were found to be capable of generating contractile forces sufficient to deform the flexible post of the Mantarray hardware after 7 days in culture (**Figure 3A; Supplementary Video 1**). Twitch responses could be achieved reliably from day 7 and forces produced from such contractions improved over time, suggesting the continued development of contractile machinery in EMTs over several weeks in culture (**Figure 3B**).

**Figure 3:**
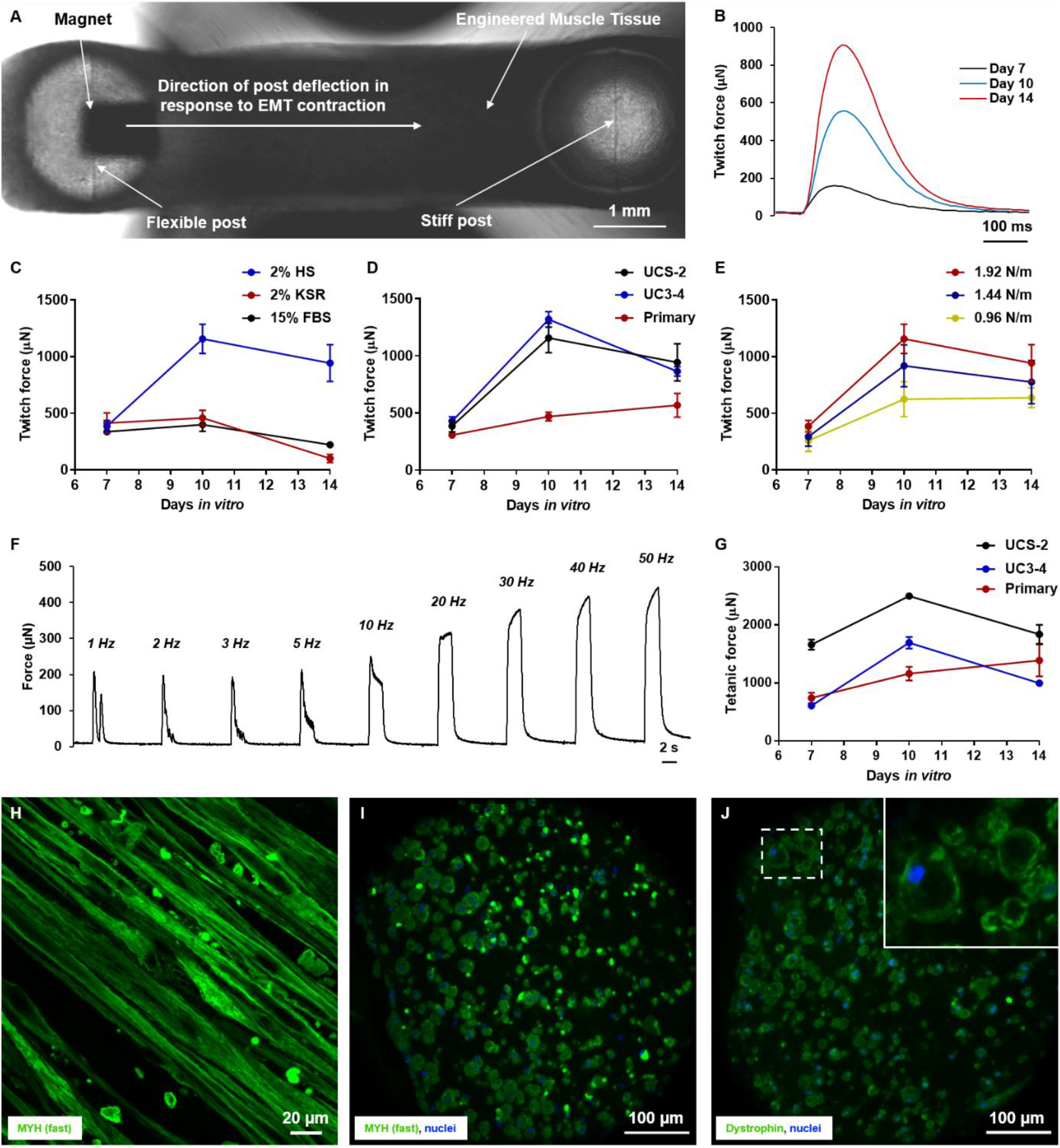
Baseline characterization of EMT function using Mantarray. (A) Representative image of a single EMT mounted in the Mantarray plate. Image was collected as a single frame from **Video 1**. (B) Representative twitch contractions from a single EMT at days 7, 10, and 14. (C) Average twitch forces measured from UCS-2 EMTs maintained in different feeding media. n = 3 to 8 per experimental group. (D) Average twitch forces measured from EMTs derived using different iPSC and primary cell lines. n = 11 to 12 per experimental group. (E) Average twitch forces measured from UCS-2 EMTs established using flexible posts of different stiffnesses. n = 3 to 8 per experimental group. (F) Representative force-frequency recording demonstrating increasing maximal force at higher frequency stimulation in a UCS-2 EMT analyzed after 7 days in culture. (G) Average tetanus forces measured from EMTs derived using different iPSC and primary cell lines and stimulated at 30 Hz. n = 11 to 12 per experimental group. (H) Representative longitudinal image of a UCS2 iPSC-derived EMT following 10 days in culture. Cells were stained with an antibody against fast myosin heavy chain prior to imaging. (I) Representative image of a UCS2 iPSC-derived EMT cross-section following 10 days in culture. Cells were stained with an antibody against fast myosin heavy chain prior to imaging. (J) Representative image of a UCS2 iPSC-derived EMT cross-section following 10 days in culture. Cells were stained with an antibody against fast dystrophin prior to imaging. Inset is a magnified image of the area outlined in white.

Given that the presence of serum in culture media represents an uncontrolled variable that can impact drug potency ^30^, many preclinical drug screening modalities identify serum-free culture conditions as preferrable. As such, we next investigated the impact of switching EMTs to serum-free medium. Fusion of iPSC-derived myoblasts to myotubes *in vitro* is achieved via standard serum starvation techniques, specifically, a reduction from 15% fetal bovine serum (FBS) to 2% horse serum (HS). In order to obtain a serum-free formulation, we replaced 2% HS with 2% knockout serum replacement (KSR) during the fusion phase of culture. In addition, we investigated whether returning EMT cultures to a high serum (15% FBS) environment, to provide additional trophic support to myotubes, following 7 days in the low serum conditions (2% HS) necessary to induce fusion had a positive effect on force development.

Of the 3 media formulations investigated, maintenance of EMTs in 2% HS throughout culture was found to promote the greatest degree of force development (**Figure 3C**). Under these conditions, average twitch force increased significantly from 385.8 µN ± 51.1 at day 7 to 1156.3 µN ± 129.1 by day 10 (day 7 versus day 10 p = 0.0007). Force production appeared to plateau at this point, measuring 943.1 µN ± 162.1 at day 14 (day 10 versus day 14 p > 0.99), with no discernable improvement thereafter (data not shown). Treatment of EMTs with 2% KSR to induce fusion produced notably poorer force development. Average twitch forces measured at days 7, 10, and 14 were 413.0 µN ± 90.3, 459.3 µN ± 66.2, and 100.2 µN ± 37.2, respectively. Similarly, treatment with 15% FBS after 7 days in culture produced average twitch forces of 337.4 µN ± 26.0, 399.6 µN ± 59.5, and 321.7 µN ± 11.4 on days 7, 10, and 14, respectively. No significant differences in the twitch profiles of UCS-2 and UC3-4 derived EMTs were observed at days 7, 10, or 14 (**Figure 3D**), highlighting the consistency of the functional profiles developed using the described methods.

Using 2% HS, we then investigated whether maintaining EMTs on post arrays incorporating flexible posts of softer stiffnesses affected force development (**Figure 3E**). While 5 different post stiffnesses were fabricated, posts with stiffnesses of 0.16 N/m and 0.48 N/m bent under passive tension to such a degree that no further deflection under active contraction could be observed. However, stiffnesses of 1.92 N/m (as used in previous experiments), 1.44 N/m and 0.96 N/m facilitated successful data capture. The collected data demonstrated that softer posts promoted the generation of smaller forces, suggesting that increasing resistance to contraction helps promote skeletal muscle function as is observed *in vivo* ^31, 32^. At day 10 specifically, posts with stiffnesses of 1.92, 1.44, and 0.96 N/m produced average twitch responses of 1156.3 µN ± 129.1, 919.7 µN ± 185.3, and 624.0 µN ± 154.6, respectively (1.92 N/m versus 0.96 N/m p = 0.04). Based on the lack of further reproducible increases in functional output after day 10, subsequent analysis of iPSC-derived EMTs were all performed at this time-point.

In addition to measuring twitch forces, higher frequency stimulation was found to be capable of driving EMTs into tetanic contractions. Longitudinal measurement of EMT contractions demonstrated that cultured iPSC-derived EMTs exhibited stepwise increases in force production in response to stimulation at increasing frequencies (**Figure 3F**), mirroring behavior observed *in vivo* ^*33*^. Tetanic force development in UCS-2 EMTs followed a similar profile as twitch function between days 7 and 14 (**Figure 3G**). Average tetanic forces measured at days 7, 10, and 14 were 1660.7 µN ± 88.7, 2498.4 µN ± 58.3, and 1839.0 µN ± 162.2, respectively. While no difference in twitch forces were observed between UCS-2 and UC3-4 EMTs, UC3-4 tissues exhibited significantly smaller tetanic output compared with their UCS-2 counterparts. Average tetanic forces for UC3-4 EMTs were 607.9 µN ± 27.5, 1693.7 µN ± 99.5, and 995.3 µN ± 54.7 at days 7, 10, and 14, respectively.

Immunocytochemical staining of day 10 EMTs highlighted the formation of striated myotubes within these constructs (**Figure 3H**). Moreover, analysis of tissue cross-sections confirmed the uniform distribution of myotubes throughout the construct as well as the appearance of dystrophin rings surrounding individual myotubes, suggesting successful attachment of the cultured cells to the surrounding matrix (**Figure 3I,J**).

### Functional characterization of EMTs derived from primary muscle cells on Mantarray

EMTs generated using primary human myoblasts produced twitch and tetanic forces that were comparable to levels observed in iPSC-derived tissues (**Figure 3D and G**). Average twitch forces measured at days 7, 10, and 14 were 306.5 µN ± 22.2, 468.8 µN ± 37.5, and 568.0 µN ± 103.7, respectively. Average tetanic forces were 740.8 µN ± 91.3, 1160.5 µN ± 118.8, and 1387.7 µN ± 275.7 at days 7, 10, and 14, respectively. While iPSC-based tissues appeared to rapidly increase force output to plateau by day 10 in culture, primary tissues displayed a comparably slower development in force progression and in certain cases, showed no change at all. One long-course study of force development found no significant improvement in twitch or tetanic force output over 30 days in culture (**Figure S2**), suggesting that primary cells produce a more stable EMT functional profile using the currently optimized culture conditions.

### Drug screening using EMTs on Mantarray

In order to verify that EMTs in culture elicited functional responses to pharmacological intervention that mirror those seen *in vivo*, we examined the impact of EMT exposure to increasing doses of 2,3-butanedione 2-monoxime (BDM) and ryanodine. BDM is a well-characterized, low-affinity, non-competitive inhibitor of skeletal muscle myosin-II ^34^ that has been shown to reversibly inhibit rat soleus muscle twitch and tetanic contractions in a dose-dependent manner without impacting action potential amplitude or time course ^35^. UCS-2 EMTs exposed to increasing doses of BDM likewise exhibited a reversable, dose-dependent inhibition of their contractile capacity, with a calculated IC_50_ of 2.5 mM (**Figure 4A,B**). Primary EMTs showed a similar response to their iPSC counterparts, with a calculated IC_50_ of 2.9 mM. This difference in calculated IC_50_ was not found to differ significantly between the experimental groups (p = 0.43 using an extra sum-of-squares F test).

**Figure 4:**
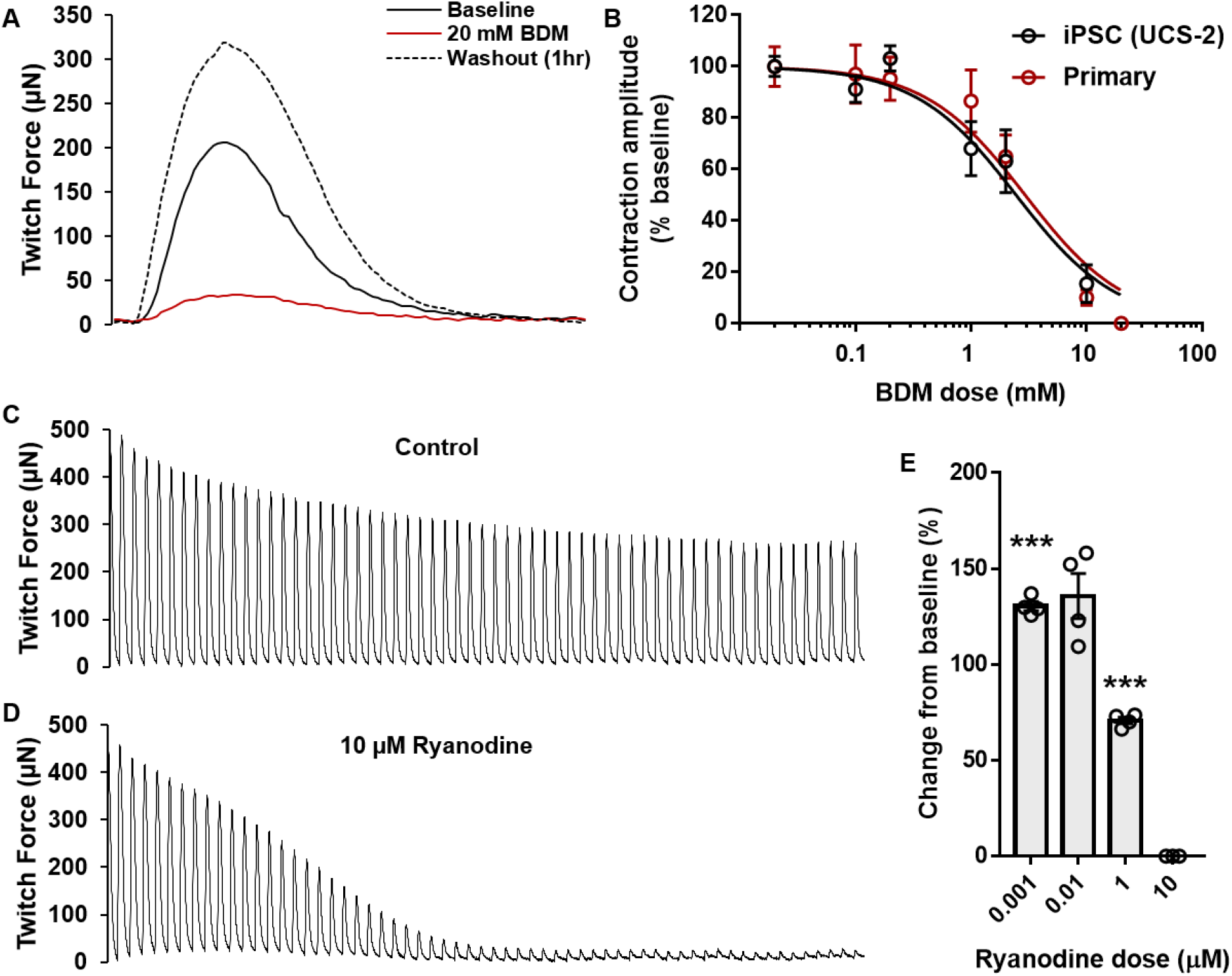
Drug responses measured on Mantarray EMTs. (A) Representative traces overlaid from a single EMT at baseline, following 20 mM BDM treatment, and after a 1-hour washout. (B) Normalized dose response curves illustrating the effect of increasing concentrations of BDM on twitch contraction magnitude in iPSC and primary muscle cell-derived EMTs. The R^2^ values for the iPSC and primary EMTs were 0.86 and 0.92, respectively. n = 4 per dose per cell type. (C,D) Representative traces from UCS-2 EMTs illustrating contraction waveforms under 1 Hz stimulation at baseline (Control) and 5 minutes after treatment with 10 µM ryanodine. (E) Change in contraction magnitude (relative to baseline) for EMTs exposed to increasing concentrations of BDM for 30 minutes. ***p < 0.001, n = 4 per dose per cell type.

Ryanodine has complex effects on the conductance and gating of ryanodine receptor channels in the muscle cell’s sarcoplasmic reticulum ^36^. Exposure to sub-micromolar concentrations causes ryanodine receptor channels to exhibit partially conducting states, which leads to an increase in sarcoplasmic reticulum Ca^2+^ permeability ^37^. Conversely, micromolar or greater concentrations produce a closed state in ryanodine receptor channels, which decreases the sarcoplasmic reticulum’s Ca^2+^ permeability ^36^. This duality was illustrated in the function of UCS-2 EMTs exposed to different concentrations of ryanodine while on the Mantarray hardware. Treatment with 1 and 10 nM ryanodine produced increases in average twitch forces of 30.4% ± 2.4 and 35.7% ± 11.7, respectively. The difference observed in the 1 nM treated group compared to baseline was statistically significant (p = 0.001). Results collected from the 10 nM treated group were not significantly different than baseline (p = 0.055) but trended towards an increase in twitch force output. Treatment with 1 µM ryanodine produced twitch contractions that were, on average, 70.8% ± 1.7 of those measured at baseline (p = 0.0004) and exposure to 10 µM ryanodine led to a complete cessation of contraction in response to electrical stimulation (**Figure 4C-E**).

### Stratification of functional phenotypes between EMTs derived from healthy and dystrophic iPSC-derived muscle on Mantarray

UC3-4 iPSC-derived EMTs bearing a dystrophin-null phenotype (a model of Duchenne’s muscular dystrophy (DMD)) exhibited significant reductions in contraction magnitudes compared with healthy (wild type) controls (**Figure 5A,B**). At day 10, wild type EMTs exhibited twitch and tetanic contraction magnitudes of 1319.9 µN ± 68.2 and 1654.1 µN ± 97.6, respectively. Dystrophic EMT forces were much smaller, measuring just 414.4 µN ± 21.9 and 267.2 µN ± 20.2 for twitch and tetanic contractions, respectively (**Figure 5C,D**). It is notable that while tetanic contraction magnitudes were larger than twitch responses for wild type cells, dystrophic EMTs produced smaller contraction magnitudes when stimulated at high tetanic frequencies than at low twitch frequencies. Measurements of contraction velocity, time to peak contraction, relaxation velocity, and time to 50% and 90% relaxation were all likewise reduced in dystrophic EMTs (**Figure 5E-I**).

**Figure 5:**
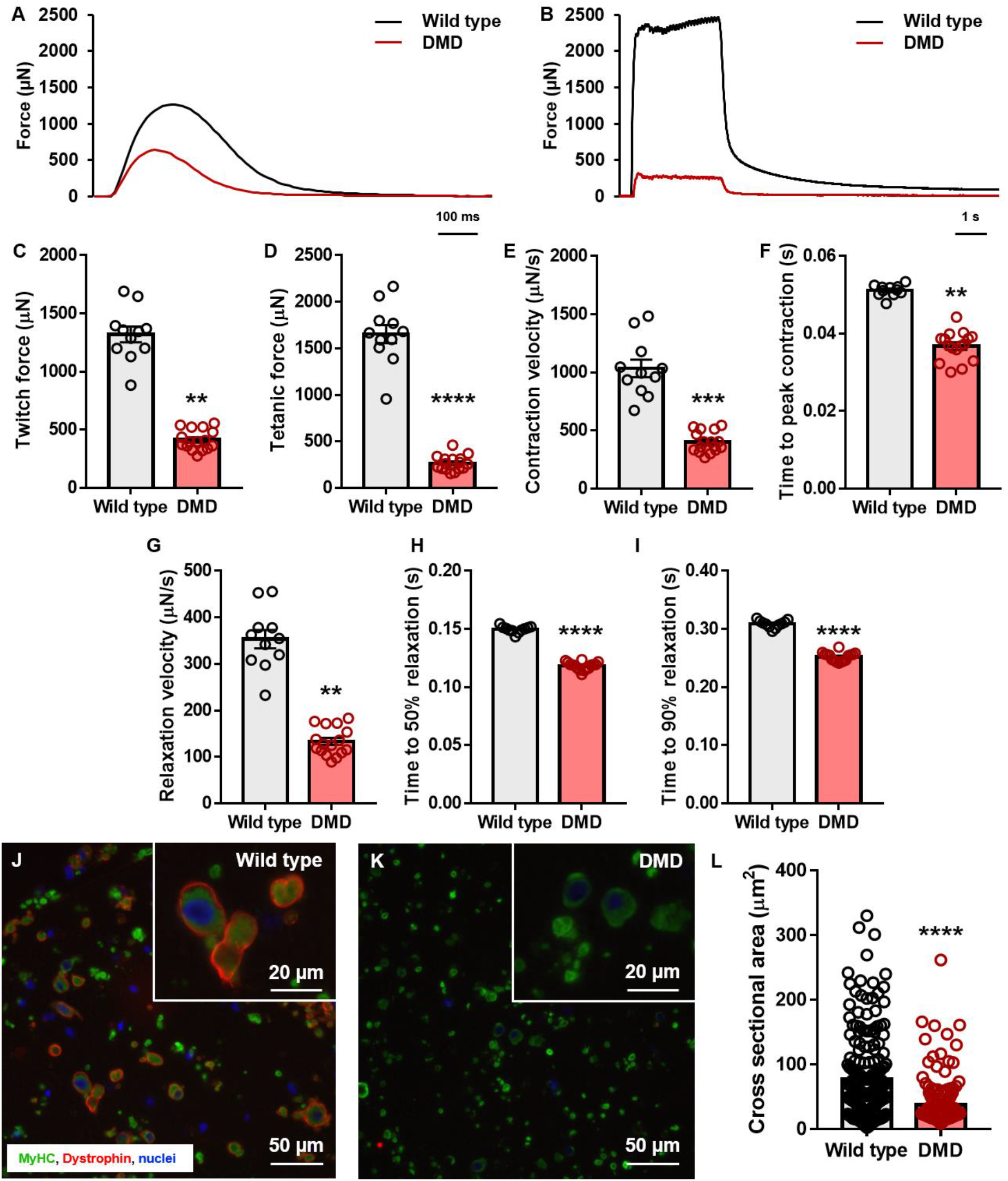
Characterizing DMD functional phenotypes in EMTs. (A) Representative twitch contractions from wild type and DMD EMTs. (B) Representative tetanic contractions from wild type and dystrophin-null (DMD) EMTs stimulated at 30 Hz. Using Mantarray, average (C) twitch force, (D) tetanic force, (E) contraction velocity, (F) time to peak contraction, (G) relaxation velocity, (H) time to 50% relaxation, and (I) time to 90% relaxation were measured across multiple tissues simultaneously. (J) Representative cross-section of a UC3-4 wild type iPSC-derived EMT following 10 days in culture. Cells were stained with antibodies against myosin heavy chain and dystrophin prior to imaging. Inset provides a magnified detail of an area from the main image. (K) Representative cross-section of a UC3-4 DMD iPSC-derived EMT following 10 days in culture. Cells were stained with antibodies against myosin heavy chain and dystrophin prior to imaging. Inset provides a magnified detail of an area from the main image. (L) Quantification of myotube cross-sectional area calculated from representative cross-sections from wild type and DMD EMTs (n = 200 per condition). For all data, *p < 0.05, **p < 0.01, ***p < 0.001, ****p < 0.0001, and n = 12 per experimental group.

To correlate observed differences in function with tissue architecture, cross-sections of both wild type and dystrophic EMTs were collected at day 10 and stained for myosin heavy chain and dystrophin expression (**Figure 5J,K**). Wild type tissues contained myosin positive cells surrounded by rings of dystrophin protein, suggesting well organized anchoring of cells to the surrounding matrix. As expected, dystrophic EMTs were found to contain myosin positive cells that completely lacked surrounding dystrophin rings. Quantification of muscle cell cross-sectional area revealed that cells in dystrophic tissues were smaller than their counterparts in control tissues (**Figure 5L**). Specifically, mean cross-sectional area in wild type and dystrophic EMTs was calculated as 76.9 µm^2^ ± 4.7 and 38.0 µm^2^ ± 2.3, respectively.

## Discussion

Despite the existence of high-throughput platforms supporting 96 engineered muscle tissues simultaneously ^38, 39^, functional assessment of these multiplexed models continues to rely on low throughput imaging techniques or sequential analysis of individual tissues attached to force transducers. Such methods reduce capacity for investigators to study the effects of prolonged (minutes to hours) stimulation on EMT function and complicate efforts to perform simultaneous assessment of multiple tissues exposed to different concentrations of drugs for accurate assessment of dose responses following defined lengths of exposure. The use of magnetic sensing as described in this manuscript enables continuous, longitudinal assessment of EMT function across 24 wells simultaneously, and higher-throughput modalities (96 and 384-well formats) are theoretically possible.

The Mantarray system enables measurements of EMT function to be performed in culture incubators, thereby further ensuring a stable recording environment that does not require mounting on microscope stages for prolonged periods. The data presented herein serves to illustrate the utility of this system for measuring the extent and reproducibility of engineered muscle performance under various conditions, including changes in cell source, post stiffness, stimulation frequency, and media formulation. In particular, we provide workflows for the production of expandable and bankable iPSC-derived myoblasts that can be stored for the establishment of large-scale screening experiments or shared between labs to help facilitate consistent results across sites. These cells can be further pushed to fuse into multinuclear myotubes that exhibit ordered sarcomere structures and contract under broad-field electrical stimulation. Incorporation of these cells into the Mantarray device and subsequent analysis of contractile output underscores that these tissues produce forces in comparable ranges to those obtained from EMTs established using primary human muscle cells. Moreover, iPSC-derived EMTs in Mantarray exhibit positive force-frequency relationships after just 7 days in culture, underscoring the capacity for this model to accurately reflect the functional profiles of native skeletal muscle ^33^ following relatively short culture periods. Confirmation of the presence of striated myotubes, alined to the surrounding extracellular matrix via dystrophin-containing protein complexes provides additional confirmation of the ability for Mantarray EMTs to reflect native tissue architecture.

The ability to establish EMTs using iPSC-derived myoblasts generated without the use of forced over expression of myogenic factors such as *Pax7* ^22^ and *MyoD* ^40^ greatly increases the potential to model neuromuscular disease states *in vitro* as patient and/or CRISPR-engineered mutant iPSC lines can be employed without the need to first install transgenes capable of driving the myogenic program. When coupled with technology for producing iPSCs from patient urine samples ^24^, the potential to expand disease modeling efforts in this area is even more profound. Our data illustrate that dystrophin-null iPSC-derived myogenic cells incorporated into the Mantarray hardware exhibit a profound functional phenotype when compared with isogenic controls. Such tissues are easily amenable to subsequent pharmacological screening to identify compounds capable of ameliorating the observed functional defect. While we have focused on Duchenne muscular dystrophy as proof of concept, the platform is adaptable to the study of any disease-causing mutation that promotes defects in musculoskeletal contraction.

Despite the differences in observed in contraction magnitude between normal and dystrophic EMTs, the presented data describe a relatively consistent contraction waveform. This may indicate that the observed difference in overall force production was due more to the size and distribution of myotubes and/or their attachment to the surrounding matrix in dystrophic EMTs rather than underlying differences in the energetics and/or biomechanics of muscle fiber contraction between these groups. Quantification of muscle cell cross-sectional area confirmed that dystrophic EMTs produced smaller cells than normal controls, providing support for this supposition. Given the presence of a purified myogenic population in these cultures, the observed defects in contraction and muscle cell hypertrophy likely reflect the effect of missing dystrophin on the contraction of skeletal muscle fibers and their connection to the surrounding matrix, specifically, without the confounding effects of fibrosis and fat infiltration as is observed in DMD patients. The current model can therefore be seen as useful for studying the intrinsic functional properties of mutant myotubes. Future models, however, could seek to incorporate fibroblasts and other supporting cells in order to offer a more comprehensive model of skeletal muscle function in health and disease.

The capacity for Mantarray EMTs to recapitulate established responses to BDM, as well as the complex pharmacology of ryanodine, underscores the potential for this platform as a preclinical drug screening tool. The current cell culture parameters suggest that serum containing media formulations are optimal for maximizing force output in iPSC-derived tissues. This is not ideal for drug development workflows and highlights areas of focus for further development with this technology. However, the near perfect overlap of BDM dose response effects in primary and iPSC-derived tissues (despite differences in the rate force development and magnitude of contractions recorded between these cell models) highlights the reproducibility of this model and the confidence this engenders in terms of its ability to generate meaningful data that can help inform clinical trial design and development. It should also be noted that the current model does not prevent *post hoc* processing of EMTs for standard molecular assays, such as histology (as illustrated), as well as transcriptomic and proteomic analyses. As such, the Mantarray hardware, coupled with recent iPSC-derived skeletal muscle culture advances, constitutes a powerful tool for high-throughput functional analysis, longitudinal functional profiling, and scaled tissue production with immediate practical relevance for drug screening and disease modeling applications.

## Acknowledgements

This work was supported by NIH GM131981 (Mack/Smith) and NIH HL149566 (Geisse). The authors would like to acknowledge Samantha Bremner for her assistance with calibrating the Mantarray device during preliminary experiments.

## Contributions

A.S.T.S and S.M.L. were responsible for performing the majority of the experiments and analysis of the generated data as well as writing the manuscript. J.B.D. helped establish the iPSC-derived skeletal muscle differentiation protocol. K.G. developed the Mantarray hardware and calibrated the machine. D.L., J.W.F., and N.J.C. developed the sectioning and staining protocol described in the manuscript and performed the staining experiments that led to the presented imaging data. S.J. helped develop the protocol for seeding muscle cells in Mantarray. J.H. and J.M. were responsible for generating the CRISPR-edited dystrophic lines used in this study. L.M., B.J.B., and E.C.F. helped with developing the concept for the project and provided edits to the manuscript. N.J.S. developed the magnetic sensing technology that Mantarray is based on and contributed edits to the manuscript. N.A.G. and D.L.M. managed the overall project and helped edit the manuscript.

## Conflict of interest

A.S.T.S., N.J.S., and D.L.M. are scientific advisors for Curi Bio, Inc.; the company that commercializes the Mantarray device used in this study. S.M.L, K.G., D.L., J.W.F., N.C., B.J.B., E.C.F., and N.A.G. are employees and equity holders of Curi Bio. All other authors state that they have no conflict of interest (financial or otherwise) associated with this manuscript.

## Supplementary Figures

**Figure S1:**
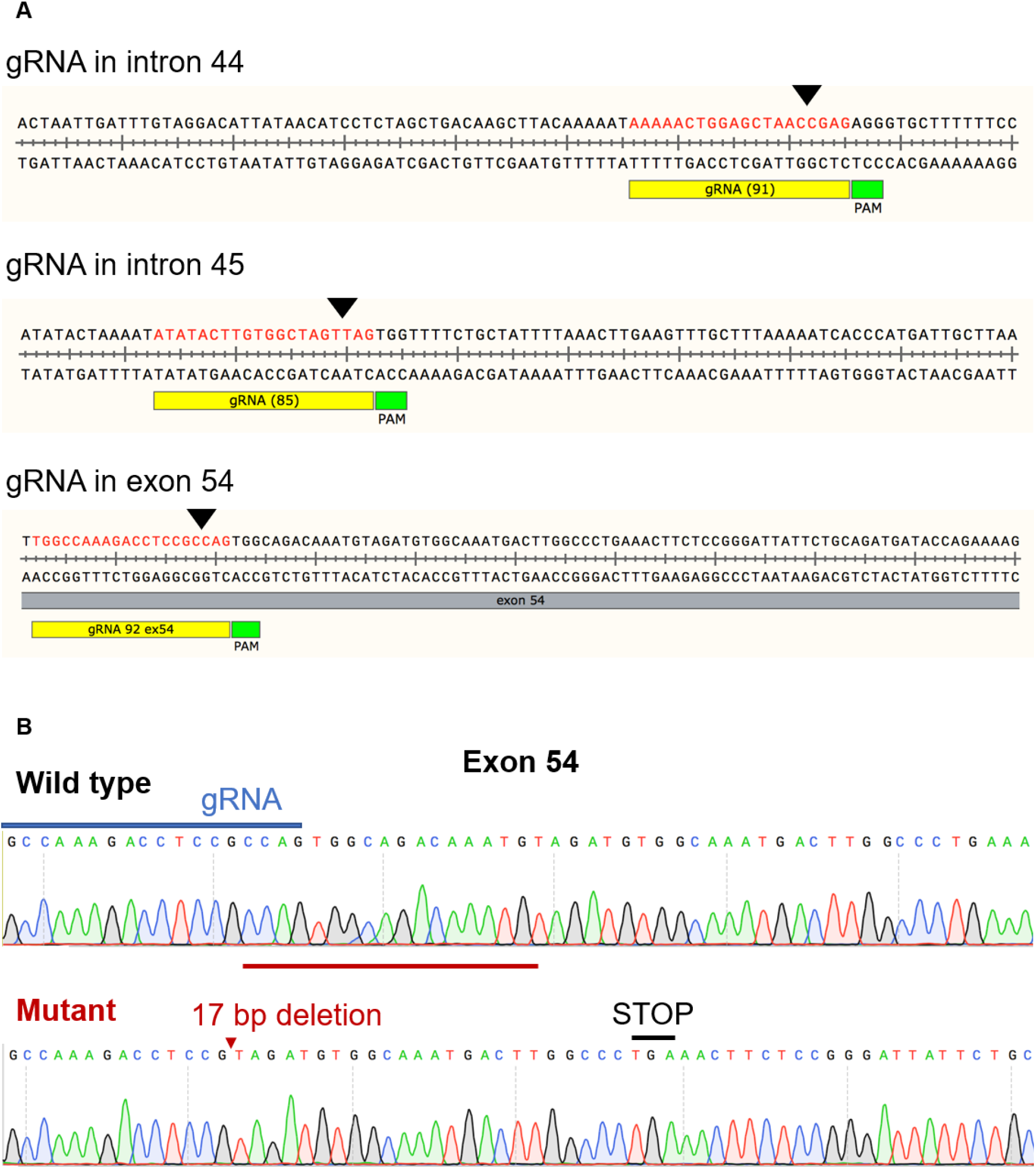
Editing strategy employed to generate the dystrophin-null line used in this study. (A) Guide RNAs used to target the dystrophin gene within the UC3-4 genotype. Black triangles indicate the cut site within each targeted intronic sequence. The blue line indicates the gRNA target sequence and the red line highlights the 17 bp sequence target for deletion. (B) Sequencing data illustrating the deletion of 17 bp in Exon 54 in order to engineer a premature stop codon. Location of the deletion is indicated by the red triangle.

**Figure S2:**
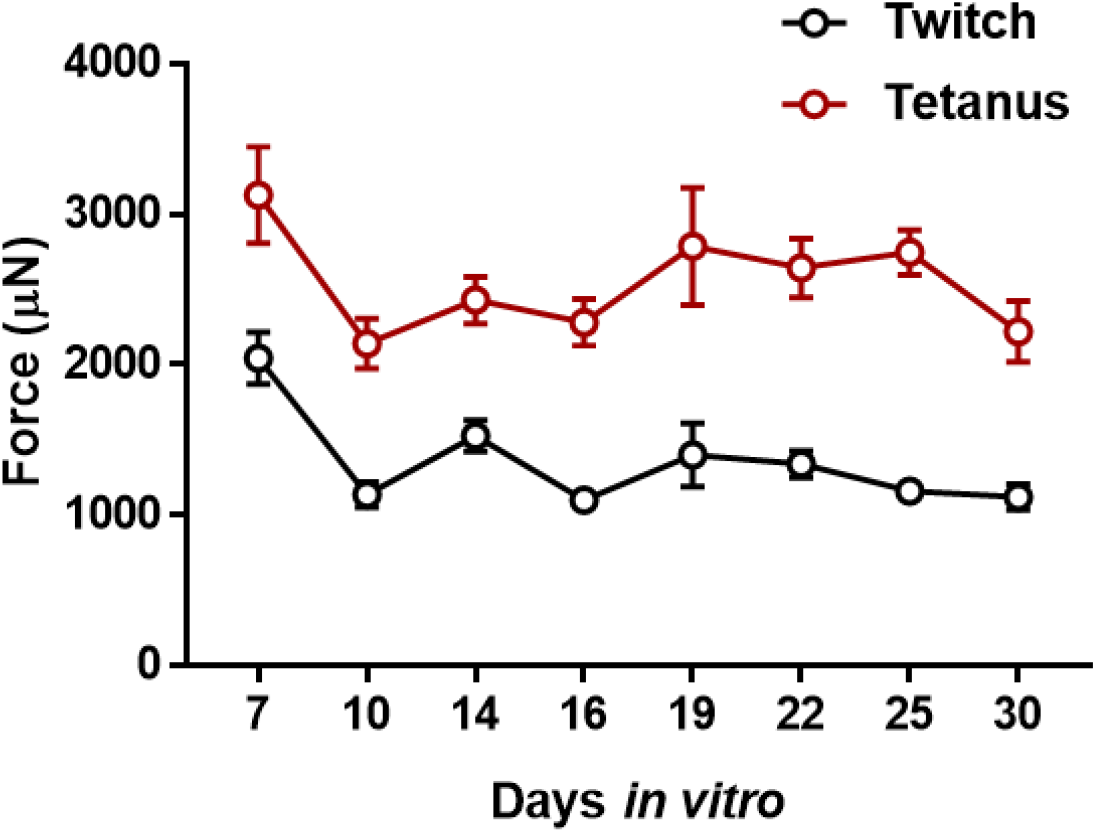
Long term evaluation of twitch and tetanic function. Primary EMTs were established and maintained in 2% HS containing medium throughout this study. n = 4.

**Supplementary Video: Representative UCS-2 EMT contracting under 1 Hz broad-field stimulation following 10 days in culture**.

## References

1. G. Agrawal, A. Aung and S. Varghese, Lab Chip, 2017, 17, 3447–3461.

2. W. Bian and N. Bursac, FASEB J, 2012, 26, 955–965.

3. K. J. Boonen, M. L. Langelaan, R. B. Polak, D. W. van der Schaft, F. P. Baaijens and M. J. Post, J Biomech, 2010, 43, 1514–1521.

4. R. G. Dennis and D. E. Dow, Tissue Eng, 2007, 13, 2395–2404.

5. R. G. Dennis and P. E. Kosnik, 2nd, In Vitro Cell Dev Biol Anim, 2000, 36, 327–335.

6. R. G. Dennis, P. E. Kosnik, 2nd, M. E. Gilbert and J. A. Faulkner, Am J Physiol Cell Physiol, 2001, 280, C288–295.

7. V. Dhawan, I. F. Lytle, D. E. Dow, Y. C. Huang and D. L. Brown, Tissue Eng, 2007, 13, 2813–2821.

8. S. Hinds, W. Bian, R. G. Dennis and N. Bursac, Biomaterials, 2011, 32, 3575–3583.

9. M. Juhas, G. C. Engelmayr, Jr., A. N. Fontanella, G. M. Palmer and N. Bursac, Proc Natl Acad Sci U S A, 2014, 111, 5508–5513.

10. A. Khodabukus and K. Baar, Tissue Engineering Part C: Methods, 2011, 18, 349–357.

11. L. M. Larkin, J. H. Van der Meulen, R. G. Dennis and J. B. Kennedy, In Vitro Cell Dev Biol Anim, 2006, 42, 75–82.

12. L. Madden, M. Juhas, W. E. Kraus, G. A. Truskey and N. Bursac, Elife, 2015, 4, e04885–e04885.

13. N. R. Martin, S. L. Passey, D. J. Player, V. Mudera, K. Baar, L. Greensmith and M. P. Lewis, Tissue Eng Part A, 2015, 21, 2595–2604.

14. V. Mudera, A. S. Smith, M. A. Brady and M. P. Lewis, J Cell Physiol, 2010, 225, 646–653.

15. D. J. Player, N. R. Martin, S. L. Passey, A. P. Sharples, V. Mudera and M. P. Lewis, Biotechnol Lett, 2014, 36, 1113–1124.

16. H. Vandenburgh, J. Shansky, F. Benesch-Lee, V. Barbata, J. Reid, L. Thorrez, R. Valentini and G. Crawford, Muscle Nerve, 2008, 37, 438–447.

17. H. Vandenburgh, M. D. Tatto, J. Shansky, J. Lemaire, A. Chang, F. Payumo, P. Lee, A. Goodyear and L. Raven, Human gene therapy, 1996, 7, 2195–2200.

18. M. R. Weist, M. S. Wellington, J. E. Bermudez, T. Y. Kostrominova, C. L. Mendias, E. M. Arruda and L. M. Larkin, J Tissue Eng Regen Med, 2013, 7, 562–571.

19. K. Yamamoto, N. Yamaoka, Y. Imaizumi, T. Nagashima, T. Furutani, T. Ito, Y. Okada, H. Honda and K. Shimizu, Lab Chip, 2021, 21, 1897–1907.

20. M. S. Sakar, D. Neal, T. Boudou, M. A. Borochin, Y. Li, R. Weiss, R. D. Kamm, C. S. Chen and H. H. Asada, Lab Chip, 2012, 12, 4976–4985.

21. O. F. Vila, S. G. M. Uzel, S. P. Ma, D. Williams, J. Pak, R. D. Kamm and G. Vunjak-Novakovic, Theranostics, 2019, 9, 1232–1246.

22. L. Rao, Y. Qian, A. Khodabukus, T. Ribar and N. Bursac, Nat Commun, 2018, 9, 126.

23. K. S. Bielawski, A. Leonard, S. Bhandari, C. E. Murry and N. J. Sniadecki, Tissue Eng Part C Methods, 2016, 22, 932–940.

24. X. Guan, D. L. Mack, C. M. Moreno, J. L. Strande, J. Mathieu, Y. Shi, C. D. Markert, Z. Wang, G. Liu, M. W. Lawlor, E. C. Moorefield, T. N. Jones, J. A. Fugate, M. E. Furth, C. E. Murry, H. Ruohola-Baker, Y. Zhang, L. F. Santana and M. K. Childers, Stem Cell Res, 2014, 12, 467–480.

25. A. S. T. Smith, C. Chun, J. Hesson, J. Mathieu, P. N. Valdmanis, D. L. Mack, B.-O. Choi, D.-H. Kim and M. Bothwell, Frontiers in cell and developmental biology, 2021, 9, 728707–728707.

26. A. S. T. Smith, J. H. Kim, C. Chun, A. Gharai, H. W. Moon, E. Y. Kim, S. H. Nam, N. Ha, J. Y. Song, K. W. Chung, H. M. Doo, J. Hesson, J. Mathieu, M. Bothwell, B.-O. Choi and D.-H. Kim, Advanced Biology, 2021, n/a, 2101308.

27. J. Chal, Z. Al Tanoury, M. Hestin, B. Gobert, S. Aivio, A. Hick, T. Cherrier, A. P. Nesmith, K. K. Parker and O. Pourquie, Nat Protoc, 2016, 11, 1833–1850.

28. J. Chal, M. Oginuma, Z. Al Tanoury, B. Gobert, O. Sumara, A. Hick, F. Bousson, Y. Zidouni, C. Mursch, P. Moncuquet, O. Tassy, S. Vincent, A. Miyanari, A. Bera, J.-M. Garnier, G. Guevara, M. Hestin, L. Kennedy, S. Hayashi, B. Drayton, T. Cherrier, B. Gayraud-Morel, E. Gussoni, F. Relaix, S. Tajbakhsh and O. Pourquie, Nat Biotech, 2015, 33, 962–969.

29. M. R. Hicks, J. Hiserodt, K. Paras, W. Fujiwara, A. Eskin, M. Jan, H. Xi, C. S. Young, D. Evseenko, S. F. Nelson, M. J. Spencer, B. V. Handel and A. D. Pyle, Nat Biotech, 2018, 20, 46–57.

30. E. Postnikova, Y. Cong, L. E. DeWald, J. Dyall, S. Yu, H. Zhou, R. Gross, J. Logue, Y. Cai and N. Deiuliis, PLoS One, 2018, 13, e0194880.

31. P. Aagaard, E. B. Simonsen, J. L. Andersen, P. Magnusson and P. Dyhre-Poulsen, J Appl Physiol, 2002, 93, 1318–1326.

32. G. R. Marcotte, D. W. D. West and K. Baar, Calcified Tissue International, 2015, 96, 196–210.

33. A. J. Fuglevand, V. G. Macefield and B. Bigland-Ritchie, J Neurophysiol, 1999, 81, 1718–1729.

34. E. M. Ostap, Journal of Muscle Research & Cell Motility, 2002, 23, 305–308.

35. M. W. Fryer, P. W. Gage, I. R. Neering, A. F. Dulhunty and G. D. Lamh, Pflügers Archiv, 1988, 411, 76–79.

36. J. L. Sutko, J. A. Airey, W. Welch and L. Ruest, Pharmacological Reviews, 1997, 49, 53–98.

37. I. N. Pessah and I. Zimanyi, Molecular pharmacology, 1991, 39, 679–689.

38. H. Vandenburgh, J. Shansky, F. Benesch-Lee, K. Skelly, J. M. Spinazzola, Y. Saponjian and B. S. Tseng, FASEB J, 2009, 23, 3325–3334.

39. M. E. Afshar, H. Y. Abraha, M. A. Bakooshli, S. Davoudi, N. Thavandiran, K. Tung, H. Ahn, H. J. Ginsberg, P. W. Zandstra and P. M. Gilbert, Sci Rep, 2020, 10, 6918.

40. A. Tanaka, K. Woltjen, K. Miyake, A. Hotta, M. Ikeya, T. Yamamoto, T. Nishino, E. Shoji, A. Sehara-Fujisawa and Y. Manabe, PloS one, 2013, 8, e61540.

